# Glycoprofiling of Cartilage Matrix-Forming Tumours

**DOI:** 10.1101/2023.01.03.522552

**Authors:** John McClure, Sheena F McClure

## Abstract

Lectin staining of benign and malignant cartilage proliferations indicates restricted glycoprofiles and significant differences. In general terms benign lesions show less cellular binding and more matrical binding than malignant lesions. This suggests that in the benign category cells are less metabolically active and the matrix more structurally stable. Chondrosarcoma cells stain with the lectin lPHA indicating increased β1-6 branching linkages in complex N-linked glycans which is also a known feature of carcinoma cells. Increased angiogenesis and increased mast cell numbers are present in the connective tissue septa separating lobules of chondrosarcoma. The blood vessels stain with the lectins PTL-II and lPHA. Mast cells stain with lPHA.

β 1-6 linkage is initiated by the Golgi-bound glycosyltransferase GnTase V. Increased β1-6 linkages are believed to enhance the metastatic potential of a malignant cell by angiogenesis. LPHA ligand is present not only in chondrosarcoma cells but also in endothelial cells and mast cells of the septa.

The ligand for the lectin PTL-II is a core 1 O-linked glycans produced under the auspices of the galactosyltransferase T synthase. Production of this glycans by endothelial cells is believed to be obligatory for the formation of these cells into tubes as part of the construction of functioning blood vessels.

There are, therefore, two candidate genes in chondrosarcoma worthy of further study viz those for GnTase V and T synthase. Targeted disruption of the activities of these genes has therapeutic possibilities.

## 1. INTRODUCTION

### Diagnostic and Therapeutic Problems of Cartilage Matrix Forming Tumours

Weber (2005) discussed what was new in musculoskeletal oncology and made a number of important points as follows:

Chondrosarcoma remains a challenging tumour to diagnose and to treat.

Adequate surgical resection is the mainstay of treatment. Chemotherapy and radiotherapy have not substantially improved the survival of patients.

Little is known about genetic aberrations and molecular pathways associated with this tumour.

Better tools are necessary for diagnosis and prediction of tumour behaviour.

The best current prognostic indication is tumour grade (Evans, 1977) which is based on light microscopy and which is highly subjective. Grading systems are not standardised between institutions. Two inter-observer studies (Eefting *et al*. 2009; SLICED Study Group 2007) have raised concerns about the high variability in diagnosis and grading of chondrosarcoma. Therefore, a more sophisticated molecular understanding of chondrosarcoma is needed.

Even low grade tumours can recur locally, change to a higher grade lesion and metastasise.

Little has changed in the intervening period since Weber made his observations and this remains the best available statement of the issues concerning chondroid (cartilage forming) neoplasms. In addition to the very important point about the capacity of seemingly lower grade lesions to transform into more malignant lesions and to metastasise, very little is known about the mechanisms of metastases of these tumours. Chondrosarcoma is a significant neoplasm since it forms 25% of primary malignant bone tumours and is the most common such tumour in the over 50 age group.

In addition to these diagnostic and predictive difficulties there are significant therapeutic challenges. Adequate surgical resection can be very mutilating and in some body sites (e.g. pelvis) technically very difficult if not impossible.

### Glycosylation and the Glycome

This project is about glycosylation which is essentially the addition of chains of saccharides (sugar molecules) to proteins. The importance of glycosylation has only relatively recently been recognised and has to do with achieving diversity after the translation of the human genome. This latter encodes for 30,000 – 40,000 proteins which is a comparatively small number and means that post-translational events must be important in achieving diversity of form, function and response to disease. The process of protein glycosylation is the most significant post translational event. Glycosylated proteins are ubiquitous components of cellular surfaces and extracellular matrices and their oligosaccharide moieties are implicated in a wide range of cell-cell and cell-matrix recognition events.

Glycosylation produces glycoconjugates (glycans) which become super informational molecules. DNA nucleotides (4) produce 256 four unit structures (<u>genome</u>). Amino acids (20) produce 160,000 four unit structures (<u>proteome</u>) and glycans can form up to 15 million four unit structures (<u>glycome</u>). That the informational content can be very high is underscored by the facts that each pair of monosaccharide residues can link in several different ways, one residue can link with 3-4 others in a branching format and glycans bearing biological information range from a few up to 20 residues. The post-translational array of glycoconjugates can usefully be termed the glycome.

For certain cancers e.g. prostate, colorectal, breast, lung and kidney there is evidence that malignant transformation, progression and metastasis are related to predictable changes in cell glycome (Pinho & Reis, 2015).

### Lectin Histochemistry

The technique of **lectin histochemistry** may be used to localise carbohydrate residues (glycans) on glycoproteins, glycolipids and glycosaminoglycans, in cell and tissue preparations.

Lectins (from the Latin legere = to select) are naturally occurring proteins and glycoproteins which selectively bind non-covalently to carbohydrate residues and which have at least two binding sites. It is for this reason that they are used histochemically (with an appropriate revealing system) as specific probes to localise defined monosaccharides and oligosaccharides from the heterogeneous mixture of carbohydrate residues on or in cells and the extracellular matrix. The Nomenclature Committee of the International Union of Biochemistry has defined a lectin as “a carbohydrate-binding protein of non-immune origin that agglutinates cells and/or precipitates polysaccharides or glycoconjugates”.

The majority of lectins have molecular weights between 60 and 130 kDa and are glycolipids. They are frequently globular proteins being freely soluble in aqueous buffers in the pH range 6-8. Lectins normally consist of two or four subunits, even through the number may be up to 20. Usually the subunits are identical and each of them has a sugar-binding site with the same specificity. The subunits are formed out of a single or of two (αβ) polypeptide chains. Isolectins are lectins which originate from the same species but have totally different specificities. PHA is a good example: E4 (ePHA) is a haemagglutinin while L4 (lPHA) causes agglutination of white cells and is mitogenic.

It is becoming apparent that carbohydrates are of fundamental importance in a range of cell recognition events in biological mechanisms as diverse as pollination in plants, fertilisation in animals, inflammation, infection and in diseases such as cancer and rheumatoid arthritis. Lectins form a large, diverse group of naturally occurring, relatively cheap, specific sugar binding molecules and are obvious tools for carbohydrate research. There are a few available good antibodies directed against carbohydrates. In contrast, there are over 100 purified lectins commercially available. By combining lectins into panels of probes detailed maps of the glycans composition of cell and tissue structures and matrices can be built up.

Lectins are very useful tools for observing glycosylation changes associated with cell development, structure, behaviour and disease. They have appeal for the biologist (relating change to cell function), for the histologist/cytologist (to distinguish cell populations), for the biochemist (in purifying glycoconjugates) and for the pathologist (detecting and understanding changes associated with disease).

### Lectins used in this study and their specificities

There have only been a limited number of studies applying lectin histochemistry to cartilage tissue. Most of these have been on animal tissue. The number (and, therefore, range) of lectins used has been small. This means that the amount of information becoming available about chemical structures and binding sites is necessarily very limited. A pioneering study by Lyons *et al*. (2007) on human knee joint cartilage developed the concept of using a panel of lectins chosen from the range of lectin groups. This maximises the amount and quality of chemical information abstracted.

Despite extensive literature searches no lectin histochemical studies of human cartilage matrix-forming tumours were discovered at the outset of this project. Given that in human articular cartilage an extended lectin panel provided a large amount of specific information and given that the matrix formed by cartilage tumours at least microscopically resembles that of articular cartilage, a panel, very nearly (but not exactly) identical to that of Lyons *et al*. (2007) was used in the present study.

Despite the obvious diversity of lectins they can, nevertheless, be classified in a number of groups depending on the nature of the chemical structures which they detect. The following is a listing of the lectins used in this study showing group, origin of the lectin and the carbohydrate specificity.

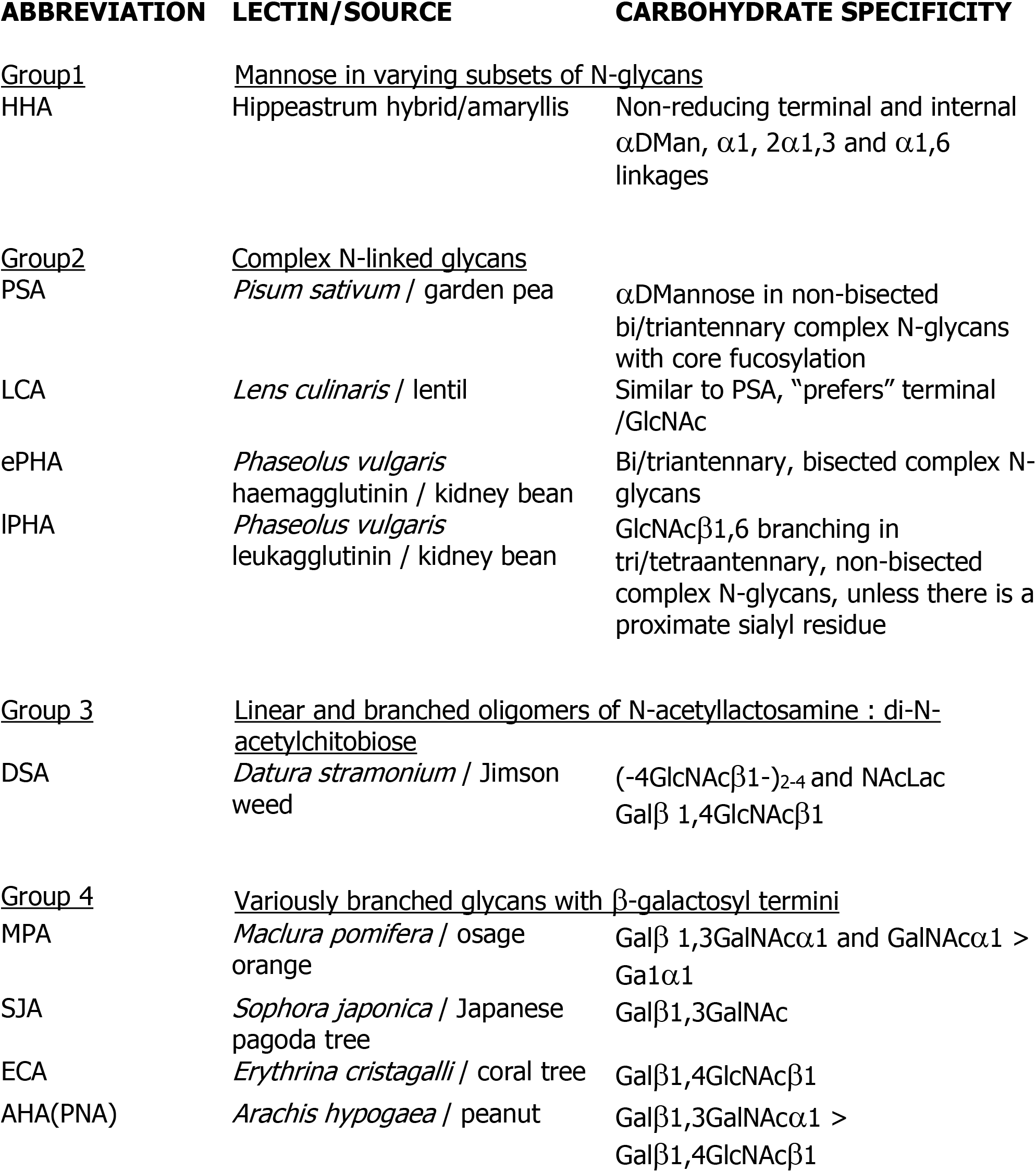

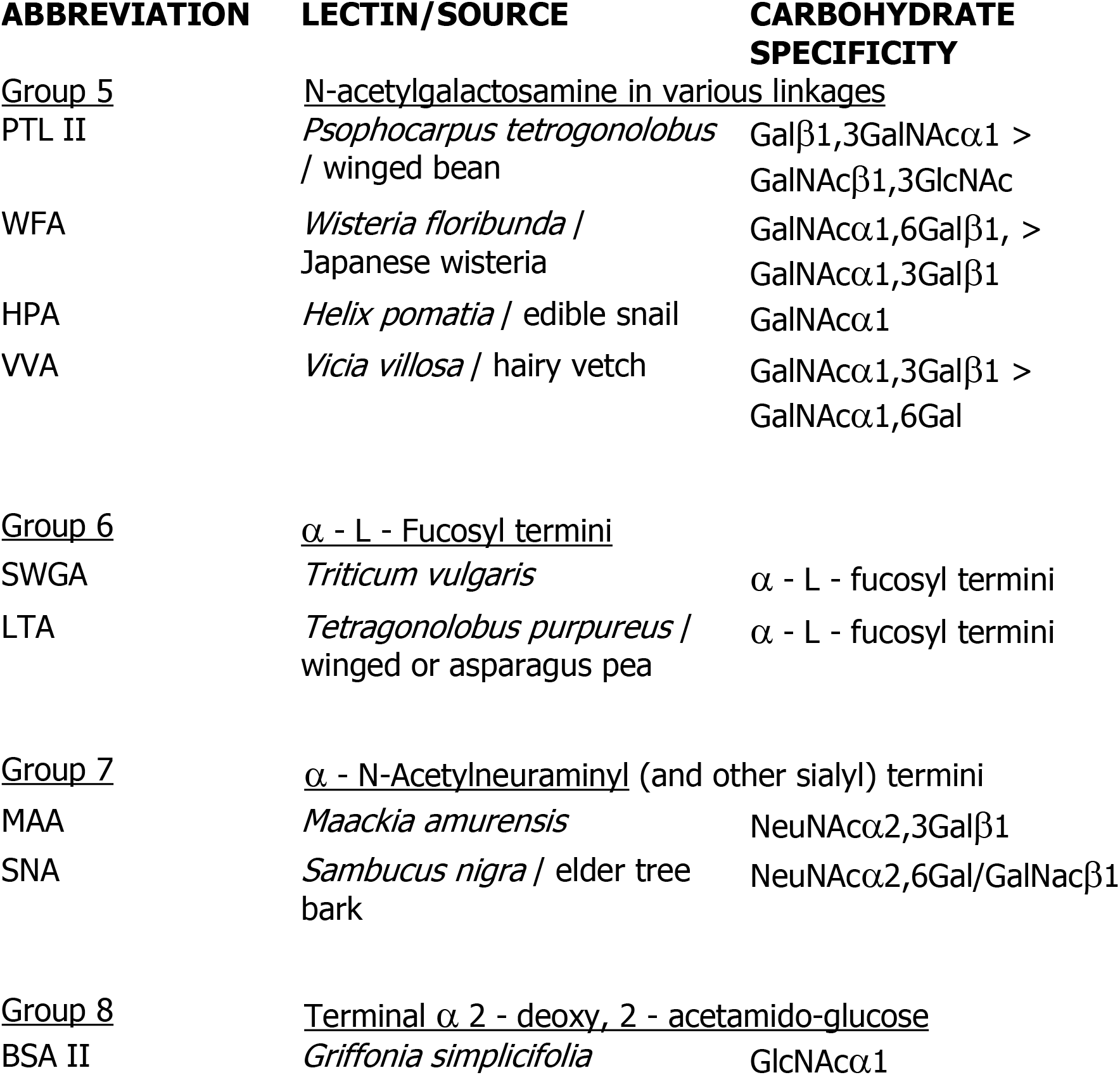

## OBJECTIVES OF THE STUDY

The objectives of the study were as follows:

To determine the glycome by examining the glycosylation pattern (i.e. glycoprofiling) of benign and malignant cartilage matrix-forming lesions.
To study the constituent cells (chondrocytes), matrix and delimiting lobular membrane of these tumours.
To determine if there are predictable differences between benign and malignant lesions.

## EXPERIMENTAL DESIGNS

### Materials and methods

A histopathology tissue archive at the onset of the study contained tissue blocks of 55 cases of enchondroma and 58 cases of conventional chondrosarcoma.

Blocks from all these cases were retrieved from the archive and new sections were cut and stained by haematoxylin and eosin (H & E). These slides were examined to provide a confirmation of diagnosis and a morphological baseline and to determine overall suitability for glycoprofiling.

In addition to the conventional H & E staining, sections containing cartilage matrix were stained by toluidine blue, picrosirius red and alcian blue in three critical electrolyte concentrations (CEC). These provided useful baseline information about the collagen and proteoglycan distribution in the matrices.

Sections were cut and stained with the panel of 19 lectins with appropriate controls. Details of the staining processes are given in Lyons *et al* 2007. Sections were also stained immunohistologically for mast cell tryptase.

All cases had been anonymised *ab initio* by one person and given a computer-generated random unique identifier number. All blocks were tested and examined without knowledge of patient demographic details and without knowledge of diagnosis. All slides were examined by another. The components of each sample were systematically examined for the presence of positive lectin staining in chondrocytes (nucleus, cytoplasm and cell membrane), matrix (pericellular, territorial and interterritorial distributions) and septal membranes (cells and blood vessels).

## RESULTS

For each case the presence of positive staining for each lectin was recorded for each micro-anatomical feature. Positive reactions were crisp, definite brown depositions. No attempt was made to grade the intensity of the reaction. This generated a large data set and in order to provide a meaningful view of the results the sum of positive reactions at particular sites was calculated as a percentage of the total number of cases per disease category. These values were shown in a compilation with lectin versus micro-anatomical site to provide an aggregated glycoprofile.

It emerged during the study that there was particular importance in the delimiting membrane and this was treated separately.

No nuclear staining was observed in any of the cases examined. This feature is, therefore, not further considered in the results presentation.

For convenience the matrix was considered in three zones – one immediately around the cell (pericellular), the second in the proximity of a cell (territorial) and the third between cells (interterritorial).

### Chondrosarcoma

All chondrosarcomas were graded on haematoxylin and eosin (H & E) stained sections by the technique of Evans *et al* (1977). The percentage of cases in the three grades were grade 1 - 23%, grade 2 - 54% and grade 3 - 23%.

Positive staining reactions were found with eleven lectins in the panel in the various micro-anatomical locations examined. No reactions were found in any of the cases in any location with the remaining lectins. The accumulated glycoprofile is shown in Table 1.

**Table 1.** Positive lectin staining reactions of chondrosarcoma. The figures shown are the percentage of cases in which staining reactions were obtained with each lectin for each micro-anatomical location. The figures in the total columns are not the sum of the figures in the adjacent columns because in individual cases two or more of the micro-anatomical features might have shown reactions.

| **CHONDROSARCOMA CELLS** |  |  |  | **MATRIX** |  |  |  |
| --- | --- | --- | --- | --- | --- | --- | --- |
| **Lectins** | **Cytoplasm** | **Membrane** | **Total** | **Pericellular** | **Territorial** | **Inter-territorial** | **Total** |
| HHA | 77 | 31 | 77 | 0 | 8 | 0 | 8 |
| PSA | 23 | 31 | 38 | 15 | 15 | 8 | 31 |
| ePHA | 62 | 15 | 64 | 23 | 8 | 54 | 54 |
| IPHA | 85 | 54 | 85 | 0 | 0 | 8 | 8 |
| ECA | 54 | 15 | 54 | 23 | 46 | 46 | 77 |
| AHA | 46 | 31 | 54 | 38 | 38 | 23 | 54 |
| SJA | 15 | 8 | 15 | 0 | 0 | 0 | 0 |
| VVA | 46 | 23 | 54 | 46 | 46 | 23 | 54 |
| PTL-II | 46 | 23 | 46 | 0 | 8 | 23 | 31 |
| HPA | 77 | 38 | 77 | 8 | 0 | 15 | 23 |
| BSA-II | 85 | 46 | 85 | 8 | 0 | 0 | 8 |

In general, lectin staining reactions were greater in cells than in matrix and greater in the cell cytoplasm than in the cell membrane. The reactions most common in cells were for lPHA, BSA-II, HHA and HPA. The cytoplasmic staining of HHA and HPA was not related to grading being equal across the grades. Cytoplasmic staining for lPHA was much lower in grade 1 compared to grades 2 and 3. The reverse situation pertained with respect to BSA-II. Membrane staining for lPHA was much greater in grades 2 and 3 in comparison to grade 1. Membrane staining for HPA was equal in grades 1 and 2 and least in grade 3. For BSA-II membrane staining was greatest in grade 1 and fell dramatically in grades 2 and 3.

In contrast, in the matrix, the most common staining reactions were for ECA, e-PHA, AHA and VVA. Staining of the inter-territorial matrix was highest in grade 3 lesions with ePHA. With ECA staining of the territorial matrix was highest in grade 1, falling to zero in grade 3. With AHA staining was equal in grade 1 and 3 and zero in grade 2. With VVA pericellular matrix staining reactions were highest in grade 3 and indistinguishably low in grades 1 and 2 tumours. The territorial reactions were equal in grades 1 and 3 and much lower in grade 2.

### Septal Membranes

Cartilage is conventionally described as an <u>avascular</u> tissue. This is only true in the sense that cartilage matrix is not permeated by blood vessels (or nerve fibres). Therefore, chondrocytes do not have an intimate association with vessels. In fact, it would appear that cartilage matrix normally has the ability to resist penetration by blood vessels. However, as in all living tissues, chondrocytes have nutritional and metabolic needs which can only be satisfied by a blood supply. In normal articular and non-articular cartilage this is achieved by a close association with the perichondrium – a vascular connective tissue which is innervated and which envelops the cartilage. The blood vessels in the perichondrium are the source of oxygen and nutrients for chondrocytes. Obviously, beyond a certain critical distance from the vessels and depending on the presence of diffusional barriers in the matrix, the chondrocyte will become hypoxic. Therefore the relationship between cartilage and its related perichondrium is key.

Similar considerations apply to malignant cartilage matrix-forming tumours. It is noticeable that the majority of tumours has a lobulated appearance composed of rounded (spheroidal) nodules separated by connective tissue septa. Observations made during the course of this study would suggest that these septa form membranes surrounding the lobules and are a parallel with the perichondrium. On haematoxylin and eosin (H & E) staining the septal membranes are composed of a collagenous connective tissue associated with spindle cells (fibroblasts/fibrocytes). Round cells are also present and these have appearances suggestive of mast cells. Blood vessels are present in the form of slit-like spaces lined by flattened endothelium which are difficult to distinguish from spindle cells. Mast cells were specifically confirmed in the septal membranes by the application of immunohistological staining for mast cell tryptase (MCT).

Glycoprofiling of spindle and mast cells and septal membrane blood vessels greatly accentuated these latter. The accumulated glycoprofiles of these constituents are shown as Table 2.

**Table 2.** Positive lectin staining reactions. The figures shown are the percentages of cases in which the several features showed staining reactions. For the lectins not shown in the table the staining reactions were universally negative.

| **SEPTAL MEMBRANES** |  |  |  |
| --- | --- | --- | --- |
| **Lectin** | **Spindle Cells** | **Round Cells** | **Blood Vessels** |
| HHA | 0 | 23 | 8 |
| PSA | 8 | 0 | 8 |
| ePHA | 8 | 8 | 23 |
| IPHA | 8 | 46 | 69 |
| ECA | 0 | 15 | 31 |
| AHA | 0 | 8 | 0 |
| SJA | 0 | 8 | 0 |
| VVA | 0 | 15 | 0 |
| PTA-II | 0 | 15 | 85 |
| HPA | 8 | 23 | 31 |
| BSA-II | 15 | 0 | 0 |

Assuming that the spindle cells are fibroblasts/fibrocytes, then their lectin reactivity is low. The most common reaction is with BSA-II (15%) and some reactions were also observed with PSA, ePHA, lPHA and HPA. In the published literature there are no comparative data available for the lectin reactivity of these cells.

Assuming that the round cells are mast cells (confirmed by immunostaining for MCT), then their glycoprofile shows particular features. Reactions were obtained with nine lectins. lPHA was the most common reaction followed by HHA, HPA, ECA, VVA, PTL-II, ePHA, AHA and SJA (in much smaller percentages). The percentage of cases showing lPHA and HHA staining was related to tumour grade being largest in grade 1 and least in grade 3.

In the glycoprofile of the septal membrane blood vessels the most common reaction was with PTL-II (85%), followed by lPHA (69%), ECA and HPA (both 31%) and ePHA (23%). The number of cases staining with PTL-II was not affected by tumour grade.

### Benign tumours (enchondromas)

The compiled glycoprofile is shown in Table 3:

**Table 3.** Positive lectin staining reactions in enchondromas. The numbers are the percentage of cases showing staining reactions with the listed lectins for cells and the three microanatomical locations in the matrix. No nuclear staining was ever observed and the staining reactions of cells are for cytoplasm and cytomembranes combined. Lectins from the panel not listed in Table 3 gave no staining reactions in either cells or matrix.

| CHONDROCYTES |  | MATRIX |  |  |
| --- | --- | --- | --- | --- |
| Lectins |  | Pericellular | Territorial | Inter-territorial |
| PSA | 69 | 0 | 72 | 69 |
| LCA | 46 | 51 | 50 | 47 |
| ePHA | 70 | 49 | 53 | 68 |
| IPHA | 0 | 0 | 43 | 65 |
| DSA | 45 | 0 | 48 | 66 |
| MPA | 43 | 0 | 45 | 48 |
| ECA | 0 | 47 | 49 | 0 |
| WFA | 49 | 49 | 69 | 51 |
| HPA | 0 | 0 | 0 | 47 |
| VVA | 0 | 0 | 0 | 49 |
| LTA | 0 | 0 | 51 | 50 |
| MAA | 0 | 0 | 0 | 52 |
| SNA | 43 | 40 | 45 | 48 |

The delimiting membranes were of variable thickness although generally very thin. Blood vessels were in moderate quantity, well-formed and giving occasional positive staining reactions with PTL-II. Mast cells were very infrequent.

Consideration of the glycoprofile shows that it is limited both for cell and matrix staining. Overall the percentage of cases showing reactions for particular lectins was lower than for chondrosarcoma. The cellular glycoprofile was more restricted than for chondrosarcoma cells indicating that cells are not only expressing fewer lectin ligands but also a smaller range and therefore metabolically less active. For a number of lectins e.g. HPA staining was present only in the interterritorial matrix. This is the zone most remote from the cell. In fact, relatively more matrical staining (with a number of exceptions) is seen away from the cell suggesting that the glycans in these have a structural role in a more stable matrical environment than is the case with chondrosarcoma. Overall the concept that benign cells are less metabolically active in a lower turnover environment would be consistent with generally accepted ideas about benign versus malignant cells.

Strikingly lPHA staining was absent in benign cells but present in territorial and interterritorial matrix. This is the reverse of the finding for chondrosarcoma cells. In contrast ePHA ligands were expressed by both cells and matrix of benign lesions. lPHA and ePHA are isolectins and have different properties. lPHA ligand is a specific linkage (GlcNAcβ1,6 in tri/tetraantennary, non-bisected complex N-glycans). On the other hand, ePHA, although it also combines with bi/tri-antennary, bisected complex N-glycans the combination is not with a specific linkage. Over-expression of lPHA, therefore, remains a characteristic of chondrosarcoma cells.

The high expression of PSA ligands in cells and matrix of benign lesions indicates that these are bi/triantennary complex N-glycans with core fucosylation. This is not the situation in chondrosarcoma where expression is low in cells and matrix. Only the matrix of benign lesions reacts with LTA which demonstrates α-L-fucosyl termini. Therefore both core and terminal fucosylation are features of benign lesions and not of malignant ones.

Sialylation is considered to be a feature of the surface membrane changes of carcinoma cells. di-N-Acetyl-neuraminyl and other sialyl termini are demonstrated by reaction with Group 7 lectins. Chondrosarcoma did <u>not</u> express these lectin ligands. Benign lesions, including cells and matrix, stain with SNA confirming that the binding site is NeuNAcα2,6Gal/GalNAcβ1.

### Conclusions

The limited glycoprofiles of chondrosarcoma cells and matrix and of the mast cells and blood vessels in the septal membranes are summarized below. Also shown is a brief, top level statement of the groupings representing the constituent lectin ligands.

### Chondrosarcoma

#### Cells

lPHA: Complex N-linked glycans: GlcNAcβ1,6 branching tri/tetra-antennary, non-bisected complex N-glycans, unless there is a proximate sialyl residue.
BSA-II: Terminal α-2-deoxy, 2-acetamido-glucose: GlcNAcα1
HHA: Mannose in varying subjects of N-glycans: Non-reducing terminal and internal αDMan, α1,2, α1,3 and α1,6 linkages.
HPA: N-acetylgalactosamine in various linkages: GalNAcα1.

#### Matrix

ECA: Variously branched glycans with β-galactosyl termini: Galβ1,4GlcNAcβ1
ePHA: Complex N-linked glycans: Bi/triantennary, bisected complex N-glycans.
AHA: Variously branched glycans with β-galactosyl termini: Galβ1,3GalNAcα1 > Galβ1,4GlcNAcβ1
VVA: N-acetylgalactosamine in various linkages: GalNAcα1,3Galβ1 > GalNAα1,6Gal

### Septal Membranes

#### Mast cells

lPHA: Complex N-linked glycans: GlcNAcβ1,6 branching in tri/tetraantennary, non-bisected complex N-glycans, unless there is a proximate sialyl residue.
HHA: Mannose in varying subsets of N-glycans: Non-reducing terminal and internal αDMan, α1,2, α1,3 and α1,6 linkages.
HPA: N-acetylgalactosamine in varying linkages: GalNAcα1

#### Blood vessels

PTL-II: N-acetylgalactosamine in various linkages: Galβ1,3GalNAcα1 > GalNAcβ1,3GlcNAc
lPHA: Complex N-linked glycans: GlcNAcβ1,6 branching in tri/tetraantennary, non-bisected complex N-glycans, unless there is a proximate sialyl residue.

It is clear that the important lectins in the context of chondrosarcoma are lPHA and PTL-II which belong respectively to lectin Groups 1 and 5. Ligands for lPHA are expressed by chondrosarcoma cells, septal mast cells and septal blood vessels whilst PTL-II ligands are only expressed by blood vessel endothelial cells.

Group 1 lectins bind to complex N-linked glycans and where these form branching structures (antennae) lPHA detects increased β1,6GlcNAc branching and there is evidence that this increased branching is related to cancer cell metastasis (Dennis *et al*. 1987). Glycosylation is dependent on a repertoire of enzymes located in the Golgi apparatus. The enzyme responsible for the initiation of the antenna is β1,6N-acetylglucosaminyl (GlcNAc) – transferase V (also known as Gn-Tase V or MGAT5). Granovsky *et al*. (2000) produced mice deficient in MGAT5 by targeted gene mutation. These MGAT5 -/- mice lacked MGAT5 products but appeared normal. However, when breast tumour formation and metastasis was induced by the polyomavirus middle T oncogene their frequencies were considerably less in MGAT5 -/- mice compared to transgenic littermates expressing MGAT5. In addition MGAT5 glycan products stimulated membrane ruffling and phosphatidylinositol 3 kinase-protein kinase B activation, fuelling a positive feedback loop that amplified oncogene signalling and tumour growth *in vivo*. Granovsky *et al*. postulated that inhibitors of MGAT5 might be useful in the treatment of malignant disease. MGAT5 products are important in cell adhesion and signalling but their precise roles in the metastatic process are undetermined.

Since GnTase V gene expression occurs not only in carcinoma but also in chondrosarcoma this suggests that common mechanisms of malignancy are in play and that tumour angiogenesis is as an important process in chondrosarcoma as it is in carcinoma. This is reinforced by the fact that mast cells and endothelial cells express ligands for lPHA.

There is much evidence that angiogenesis is related to mast cells which accumulate in many angiogenesis-dependent situations, including tumour growth, rheumatoid arthritis, ovulation, wound healing and tissue repair. The common belief is that mast cell products are angiogenic and regulate endothelial cell proliferation and function. In addition, mast cell products such as tryptase also degrade connective tissue matrix to provide space for neovascular sprouts (Hiromatsu & Toda, 2002). Mast cells up-regulate angiogenesis during squamous epithelial carcinogenesis (Coussens *et al*. 1999).

PTL-II is not usually thought of as a marker of endothelial cells in tissue sections and it is not known if the observation is specific for chondrosarcoma or if blood vessels are marked in other tumours. Unpublished work in the authors’ laboratory showed that PTL-II bound to human umbilical vein endothelial cells. PTL-II belongs to lectin Group 5 whose members combine with N-acetylgalactosamine in various linkages. The ligand for PHL-II is Galβ1,3GalNAcα1 > GalNAcβ1,3GlcNAc. The responsible enzyme forming the core 1 O-glycans Galβ1,3GalNAcα1 is β1,3 galactosyltransferase (T synthase). Xia *et al*. (2004) found that core 1 O-glycans in wild-type mice was expressed primarily in endothelial, haematopoietic and epithelial cells during development. Gene-targeted mice lacking T synthase developed brain haemorrhage that was inevitably fatal by embryonic day 14. Histological examination showed that in T synthase deficient brains there formed a chaotic microvascular network with distorted capillary lumina and defective association of endothelial cells with pericytes and extracellular matrix. This revealed an unexpected requirement for core 1 O-glycans during angiogenesis and that the requirement had to do with proper structural organisation of the blood vessel at the microscopical level. In fact, it has been suggested (Tian & Ten Hagen, 2009) that altogether there is a conserved role for O-glycans in the molecular events governing tubulogenesis across diverse organ systems and species, possibly by influencing trafficking and stability of crucial proteins involved in tube architecture and function. For patients with chondrosarcoma, the overall prognosis is related to size of the lesion, anatomical location and histological grade. Patients with axial lesions have a worse prognosis than those with tumours of the appendicular skeleton. The five year survival rate for patients with grade 1 lesions is 90%, the rate decreases to 29% with patients with grade 3 tumours. Grade 1 lesions do not metastasise. Metastatic spread, typically pulmonary, is more frequently associated with grade 3 lesions than with other grades of lesions. Lymph node spread is more common with chondrosarcoma than with other osseous neoplasms. Tumour recurrence typically occurs 5-10 years after surgery. Recurrent chondrosarcoma is often more aggressive than the original lesion and the histological grade is often higher.

These observations underscore the apparent importance of histological grade in prognosis of outcome for chondrosarcoma. However, serious questions have recently been raised about the validity of grading. There is very considerable between-observer variation and there is no agreed method of harmonising approaches in different treatment institutions. As with all grading systems there is a more fundamental problem. Grading seeks to impose discrete variables on what is, in biological terms, a continuum. In other words, at the boundaries between grade values there are bound to be overlaps. Applying a grade value is, perhaps, no more sophisticated than saying that tumours which microscopically look benign will most likely behave in a benign way whilst those which look malignant will most likely behave in a malignant way.

The clear distinction of the glycoprofiles of enchondromas and chondrosarcoma permits separation of benign and malignant tumours. Glycoprofiling by lectin histochemistry is a lengthy process and its usefulness in acute diagnosis is doubtful. However, the moment of mass glycomics is upon us offering opportunities for timely diagnoses.

The most important findings to emerge from the present study relate to the vascularity of the delimiting septal membranes around the typical lobulated structures of chondroid tumours. New thin-walled blood vessels are characteristically stained by, in particular, PTL-II. These vessels are very difficult to visualise in H & E staining, since they lack a prominent basement membrane. Angiogenesis is a well-recognised phenomenon in common carcinomas such as breast and colon, and a lot of clinical effort has been directed at developing therapies designed to inhibit angiogenesis and there are encouraging advances in this context.

As has been stated previously it is commonly assumed that cartilage is an avascular tissue. This is true in the sense that cartilage matrix is not permeated by blood vessels and cartilage cells do not have an intimate relationship with fine capillary structures. However, the vasculature of cartilage is organised in and limited to the peripheral membrane structure in the perichondrium of normal cartilage and the septal membranes of chondroid tumours. The survival of the cell is, therefore, critically dependent on its distance from the blood vessels. In consequence the cartilage tumour lobule has a maximum size beyond which there will be central hypoxia and possible necrosis. For a cartilage tumour to proliferate and expand the density of the peripheral vasculature must also expand. The present study clearly demonstrates that this occurs. More malignant tumours recruit more blood vessels.

In addition to new blood vessels providing for the nutritional and metabolic needs of growing tumours, these vessels are believed to be incomplete structurally – lacking developed and intact basement membranes and are believed to be “leaky”. The ultimate malignant behaviour of any tumour is metastasis viz distant spread most commonly to lungs via the blood stream. The presence of leaky vessels in increased numbers represents an easy portal of entry, which would facilitate the first part of the metastatic journey.

Mast cells were also found in the septal membrane. There have been no previous reports of mast cells associated with chondrosarcoma. For other tumours, specifically carcinomas, there have been reports of these cells congregating at tumour margins. The significance of this is uncertain. There is a belief that this is part of an anti-tumour response mechanism. The alternative view is that mast cells are trophic to the tumour encouraging growth and facilitating vascular invasion.

It is now recognised that mast cells show physiological heterogeneity. The chemical constituents of connective tissue mast cells (CTMC) are different from those of cells associated with mucosal surfaces (MMC). However, it is probably the case that this heterogeneity is even more subtle and complex. This study and other work by the authors and colleagues have shown that for CTMCs there are limited glycoprofiles which vary by tissue and disease process. The implications are that their cellular enzyme repertoires also vary in a specific fashion.

The restricted glycoprofiles of both septal membrane mast cells and blood vessels offer the potential for chemical dissection with the recognition of molecular targets for anti-tumour therapies.

## Figures

**1.**
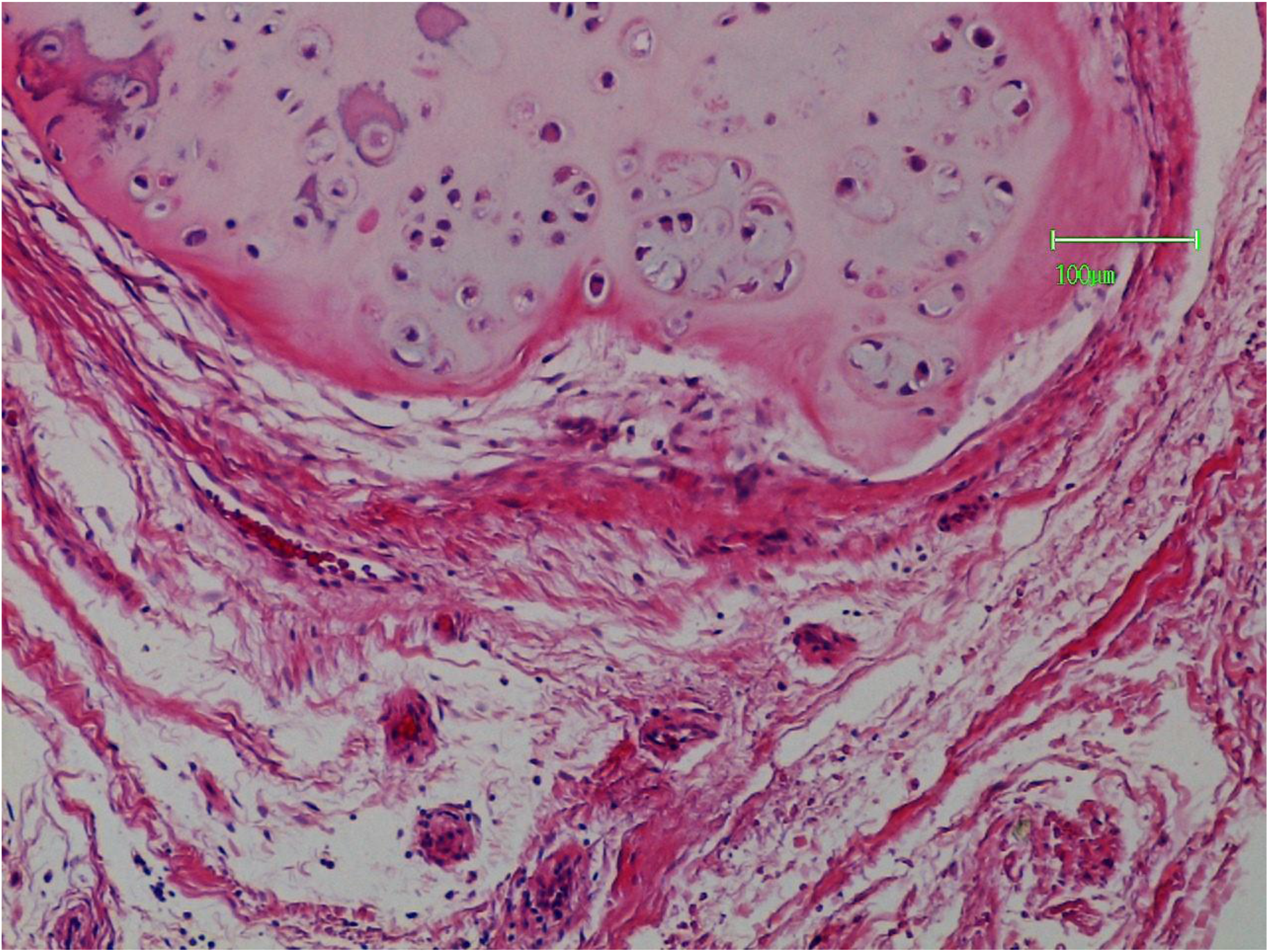
Lobule of chondrosarcoma surrounded by a vascular connective tissue containing mast cells (Haematoxylin and eosin).

**2.**
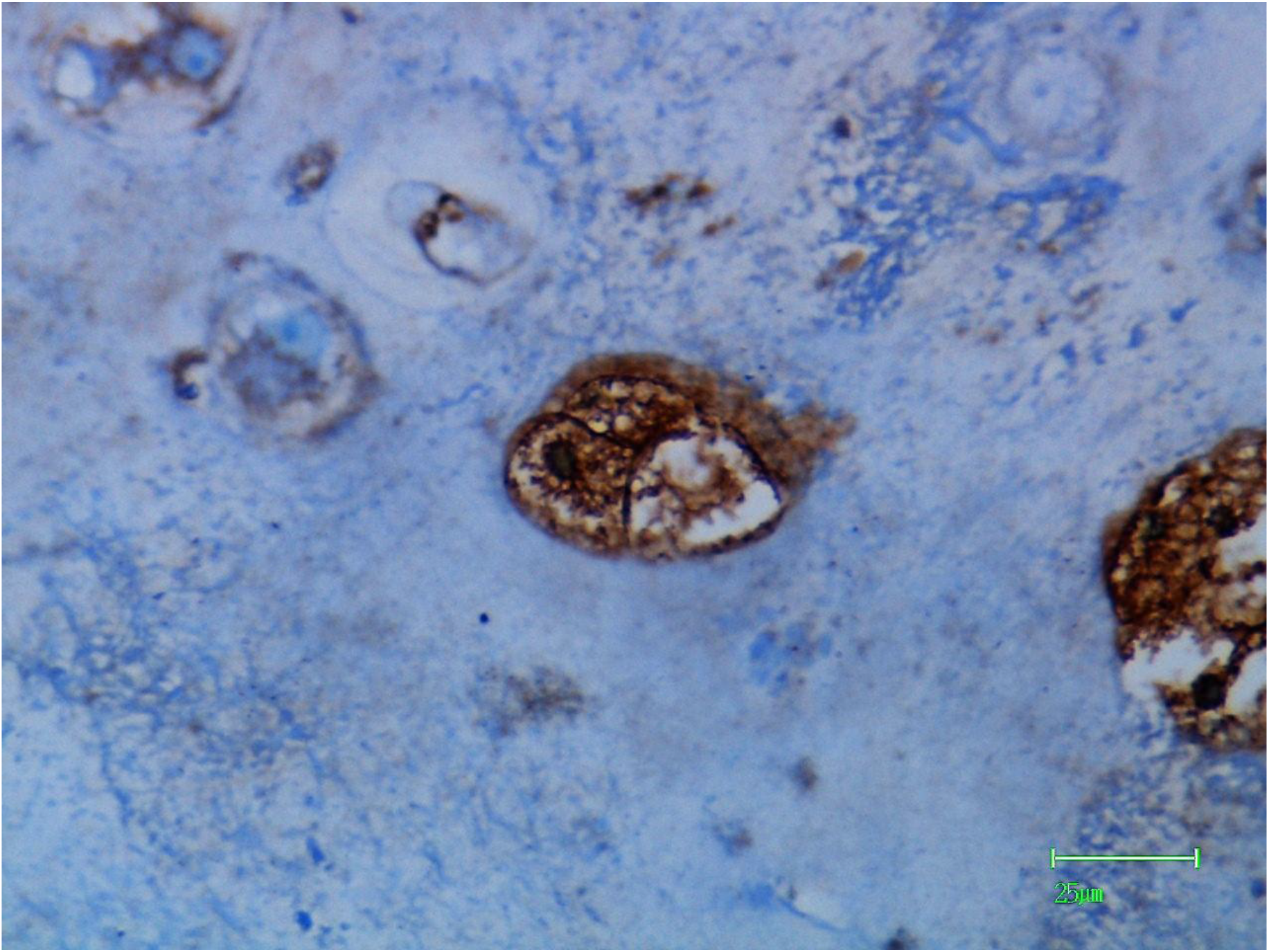
Chondrosarcoma cell expressing ligand for lPHA (brown granular material) which is cytoplasmic and membranous in distribution.

**3.**
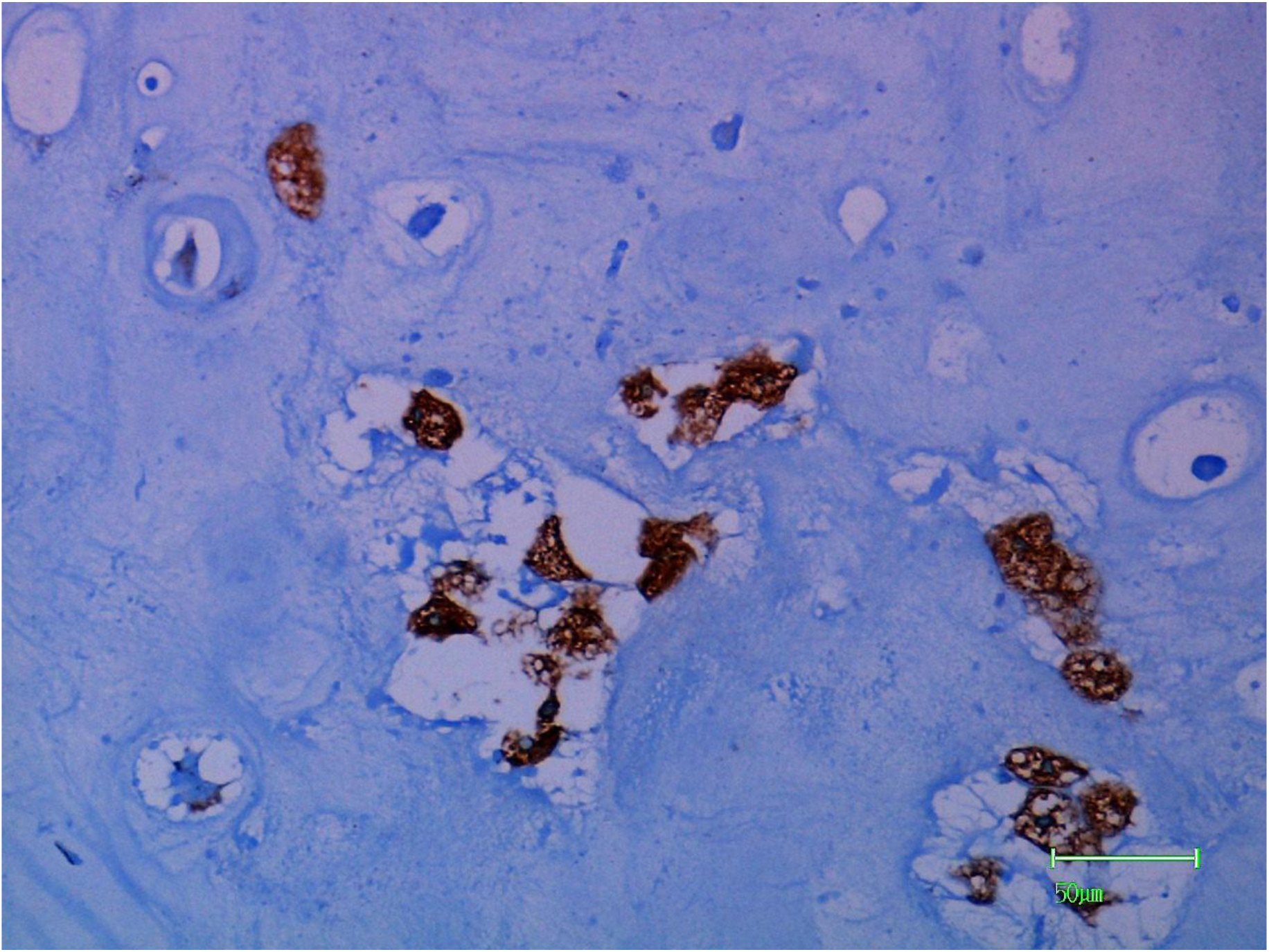
Clusters of chondrosarcoma cells showing positive reactions with lPHA. The positive material is displaced by marked intracytoplasmic hydropic change (cleared spaces).

**4.**
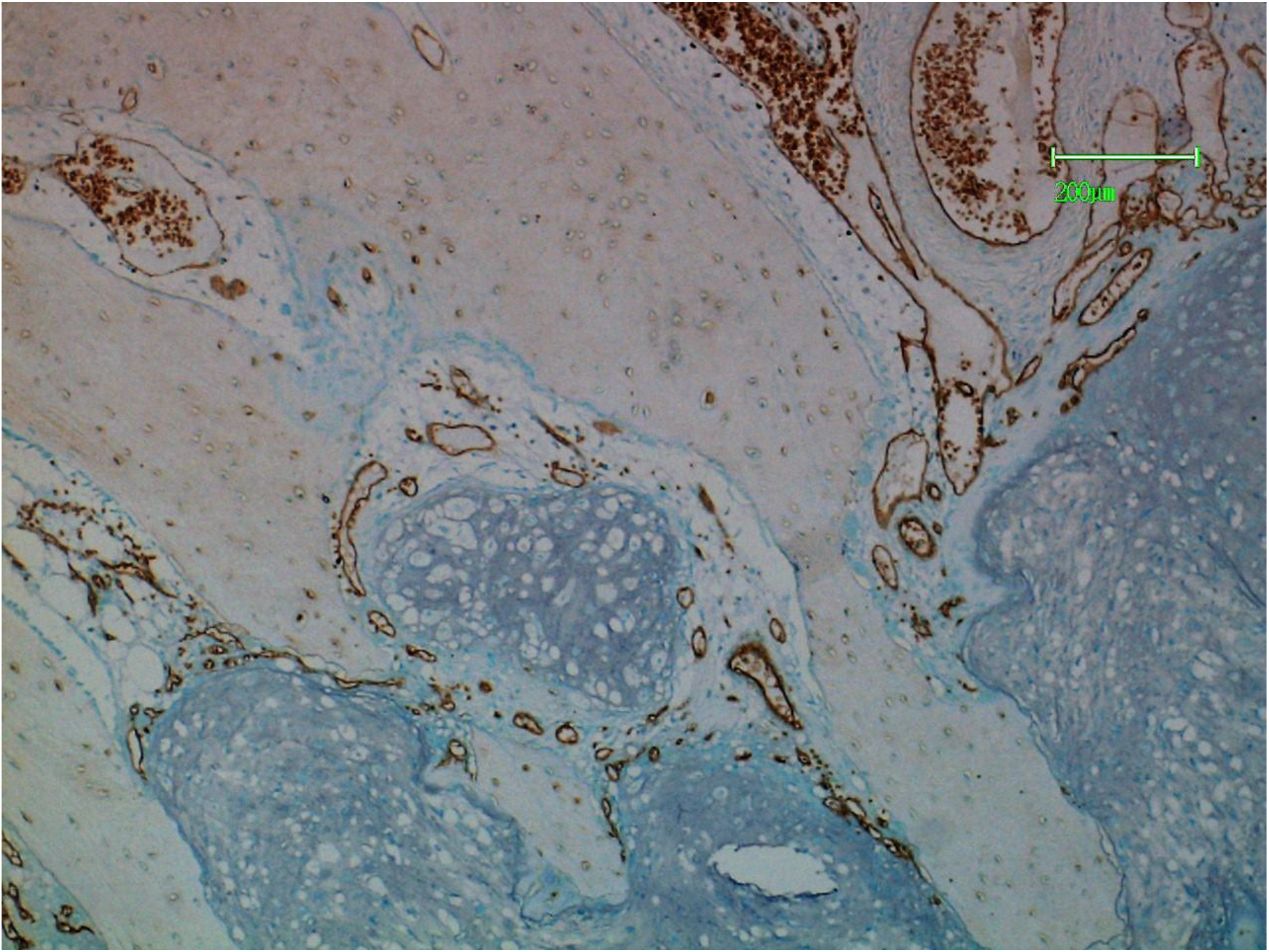
Lobules of chondrosarcoma delimited by a vascular connective tissue in which the endothelial cells stain positively with PTL-II.

**5.**
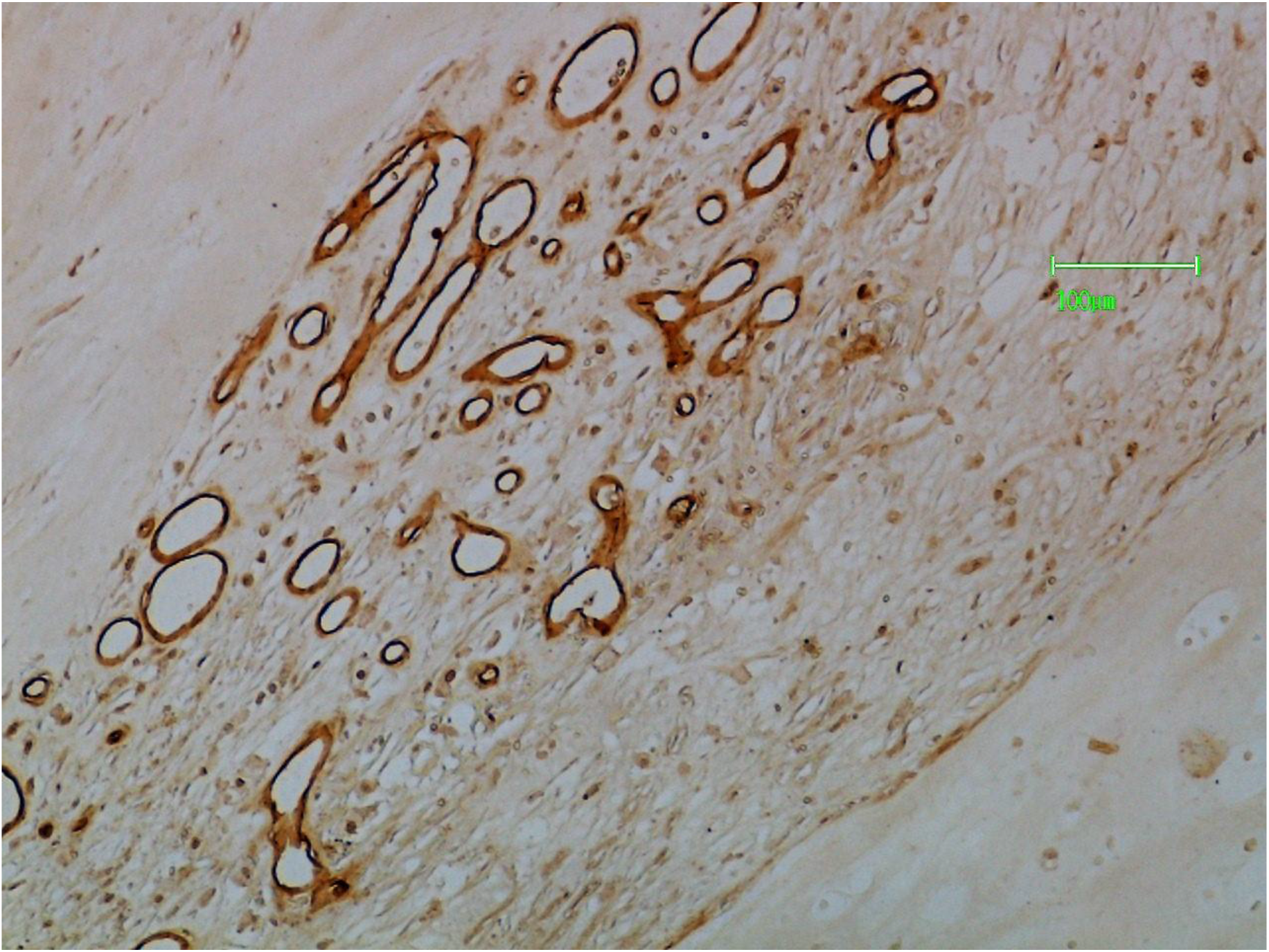
Connective tissue septa with endothelial cells staining positively with PTL-II.

